# Comparative genomics approaches accurately predict deleterious variants in plants

**DOI:** 10.1101/112318

**Authors:** Thomas J. Y. Kono, Li Lei, Ching-Hua Shih, Paul J. Hoffman, Peter L. Morrell, Justin C. Fay

## Abstract

Recent advances in genome resequencing have led to increased interest in prediction of the functional consequences of genetic variants. Variants at phylogenetically conserved sites are of particular interest, because they are more likely than variants at phylogenetically variable sites to have deleterious effects on fitness and contribute to phenotypic variation. Numerous comparative genomic approaches have been developed to predict deleterious variants, but the approaches are nearly always assessed based on their ability to identify known disease-causing mutations in humans. Determining the accuracy of deleterious variant predictions in nonhuman species is important to understanding evolution, domestication, and potentially to improving crop quality and yield. To examine our ability to predict deleterious variants in plants we generated a curated database of 2,910 *Arabidopsis thaliana* mutants with known phenotypes. We evaluated seven approaches and found that while all performed well, their relative ranking differed from prior benchmarks in humans. We conclude that deleterious mutations can be reliably predicted in *A. thaliana* and likely other plant species, but that the relative performance of various approaches does not necessarily translate from one species to another.

## INTRODUCTION

Dramatically increased numbers of reference genomes and whole genome resequencing data sets have facilitated the discovery of sequence variants and increased interest in the annotation of functional variants in many organisms. Functional annotation can yield insight into the genetic basis of phenotypic variation and is often a critical step in the identification of genes and variants underlying human disease (1, 2). In particular, interest in identifying putatively deleterious variants has increased, because these variants may contribute substantially to phenotypic variation (3, 4). Because deleterious variants are more likely to disrupt phylogenetically conserved sites, the availability of comparative genomics data has made it possible to develop computational approaches to identifying deleterious variants genome-wide (5). Although a number of approaches have been developed to identify deleterious variants within noncoding sequences (e.g. (6) and (7)), most have focused on variants that alter the amino acid sequence of proteins (5). This focus on amino acid substitutions in protein coding sequences is in part driven by the observation that amino acid-altering single nucleotide polymorphisms (SNPs) are more often associated with phenotypic variation than other classes of variants, but also because they are the most readily identifiable class of variants that are likely to have a biological impact (8–10).

While identification of disease-causing and potentially “actionable” genetic variants is fundamental to personalized medicine, identifying deleterious variants is also broadly relevant to understanding the genetic basis of phenotypic variation. In humans, annotation of deleterious variants improves prediction accuracy of complex traits (11). For domesticated organisms, complementation of recessive deleterious variants between haplotypes is thought to be one of the primary mechanisms underlying heterosis (12). This suggests that identification of deleterious alleles may be applied to hybrid breeding strategies (13). In humans, annotation of deleterious variants improves prediction accuracy of complex traits (11). Elevated proportions of deleterious relative to neutral variants in domesticated species suggest a cost of domestication (14–18). Studies of the genomic distribution and genetic contribution of deleterious variants can contribute both to understanding the origin and domestication of crop species and to advancing breeding and crop improvement strategies (19).

Accurate prediction of deleterious variants is a key component of assessing their contribution to phenotypic variation. Numerous approaches for predicting deleterious variants have been developed. The performance of an approach is typically assessed using the proportion of known, disease-causing human variants that are accurately classified as deleterious. Benchmarking of various approaches using standardized test sets has shown substantial variability among approaches, and improved performance is often achieved through combining results from multiple tools (20–23). However, the causes of performance differences across approaches are not well understood. While all approaches rely on sequence conservation at the phylogenetic level to identify deleterious variants, some approaches also incorporate protein structure, physical or biochemical properties of amino acid changes, or other attributes of protein sequence when they are available. The earliest conservation metrics used heuristic measures, sometimes including filtering or weighting to account for phylogenetic distance (24). More recent approaches have incorporated evolutionary models that account for phylogenetic distance based on putatively neutrally evolving nucleotide sites (25, 26). Reference bias and the alignments used to calculate conservation metrics are not often emphasized, but are important for making accurate predictions and may account for some of the variability among predictions (26–28). The accuracy of predictions is particularly dependent on the availability of annotated genomes among related species and the potential to generate sequence alignments.

Despite most approaches being developed for and applied to humans, there has been growing interest in identifying deleterious variants in non-human species in order to understand genomic patterns of variation and their contribution to complex traits, especially in plants. Patterns of deleterious variation have been examined in *Arabidopsis thaliana* (29), rice (15, 30), maize (17, 31), sunflower (32), poplar (33), barley, and soybean (34). However, the accuracy of predictions in plants has only been examined for a small number of known variants (30) and only in the past few years have a diverse set of plant genomes and protein homologs become available (35). Furthermore, plants are known to have a larger number of multi-gene families and a higher frequency of polyploidy than occurs in mammals (36). These genome-specific factors influence whether a sequence variant is truly deleterious in a given species (37, 38).

The goal of this study was to evaluate the ability of various approaches to predict deleterious variants in plants. The model system *A. thaliana* is a particularly attractive plant species for evaluating approaches that predict deleterious variants because decades of basic research in development, physiology, cell biology, and plant-pathogen interactions have identified large numbers of amino acid-altering mutations with phenotypic consequences. We identified seven approaches that can predict deleterious variants outside of humans (Table S1). Among these approaches, SIFT (24), PolyPhen2 (27) and PROVEAN (39) generate their own alignments using hits from non-redundant protein databases, whereas MAPP (40), GERP++ (25), and two versions of a likelihood ratio test (26) make predictions using pre-specified alignments as input (Table S1). Because new genome sequences are continually becoming available, the BAD_Mutations pipeline was developed to flexibly identify homologs and generate alignments for any protein of interest (34). BAD_Mutations uses TBLASTX (41) to identify the best match (homolog) from each specified genome and aligns them with PASTA (42). For the four approaches that require alignments, we used the BAD_Mutations pipeline applied to 42 plant genomes. BAD_Mutations was also used to implement two approaches based on a likelihood ratio test (26, 34).

To evaluate predictions of deleterious variants in plants, we generated a curated database of 2,910 *A. thaliana* mutants with known phenotypic alterations. We evaluated the ability of seven approaches to identify these deleterious variants and found that while performance was better than similar assessments in humans, the relative ranking and the highest performing approach differed from previously reported comparisons using human data. Our results demonstrate that reliable prediction of deleterious variants can be achieved in *A. thaliana*, and likely other plant species, expanding the potential value of using deleterious variants to understand naturally occurring variation and to improve crop breeding strategies.

## MATERIAL AND METHODS

We curated a set of amino acid-altering mutations with phenotypic impacts. Both morphological and biochemical phenotypes were represented, and mutations were in both single-copy and duplicated genes. These mutations were obtained from two sources. We generated a manually curated set of 542 amino acid-altering mutations in 155 genes with phenotypic effects that are described in the literature. These mutations were found by searching the *Arabidopsis* Information Resource (http://www.arabidopsis.org) for genes with either dominant or recessive alleles that differ by nucleotide substitutions. We also identified mutations using a literature search in Google Scholar (http://scholar.google.com). For each variant, we recorded the amino acid substitution, position, and link to the published paper (Table S2). We excluded nonsense mutations because they frequently completely eliminate gene function. We identified a second set of 2,617 amino acid-altering mutations in 960 genes from the manually curated UniProt/Swiss-Prot database (http://www.uniprot.org/) (43). The two sets were independently generated and had an overlap of 249 mutants. Using mutants with named alleles as a proxy for those with morphological versus biochemical phenotypes, 65% of our manually curated set and 33% of the Swiss-Prot set had macroscopic phenotypes. Duplicated genes were defined by those proteins with a significant BLASTP hit (E-value < 0.05) to another *A. thaliana* protein with > 60% identity. By this criterion 466 of 995 proteins were classified as duplicated.

Single nucleotide polymorphisms (SNPs) without any known phenotype were obtained from a set of 80 sequenced *A. thaliana* strains (Ensembl, version 81, “Cao_SNPs”, (29)). At the time of download, these were the only SNP set available for unrestricted use. After filtering out sites with heterozygous or missing genotype calls, there were 10,797 biallelic amino acid-altering SNPs in the 995 proteins. We used a subset of 1,583 common SNPs (>10%) as those least likely to have phenotypic effects. Our rationale is that on average, strongly deleterious alleles are less likely to reach high frequency in a population, owing to the effects of purifying selection (44). We also assessed performance by measuring the enrichment of deleterious variants predicted for rare compared to common polymorphisms (45). A second set of common amino acid-altering SNPs were identified in an independent set of genes. Excluding the original set of 995 genes, we randomly selected 1,000 proteins from 35,386 peptides in the *A. thaliana* database. We removed 21 that carried no amino acid polymorphism in the 1,001 genomes project (1001genomes.org). In the remaining 979 genes, we identified 40,736 biallelic amino acid altering SNPs in the 1,001 genomes project, of which 3,717 were common (>10%).

We assessed amino acid substitutions using seven approaches: LRT (26), LRT-masked (33), PolyPhen2 (46), SIFT 4G (47), Provean (39), MAPP (40) and GERP++ (25). PolyPhen2 predictions were generated using the standalone software (v2.2.2) with the PolyPhen2 bundled non-redundant database (uniref100-release 2011_12) and the probabilistic variant classifier using the default HumDiv model. Precomputed SIFT 4G predictions were obtained for *A. thaliana* (TAIR10.23) (http://sift.bii.a-star.edu.sg) and are based on the UniRef90 database (2011). SIFT 4G predictions were not available for 855 substitutions, predominantly because the amino acid change involved more than one nucleotide change within a codon. Provean predictions (v1.1.5) were generated for all mutations using NCBI's non-redundant database (04/02/2016). MAPP and GERP++ predictions were generated using BAD_Mutations alignments and trees (see below). GERP++ generates predictions for single nucleotide positions rather than codons, based on a deficit of observed substitutions compared to that expected given a neutral substitution rate. To assess GERP++ performance we used the GERP++ score at the first, second or third position of the codon if the amino acid substitution could occur by a single change at one of those positions and the average of the GERP++ scores at the first and second positions for all other types of changes. In addition, because GERP++ did not initially perform well on the *A. thaliana* data using neutral substitution rates estimated from each alignment (default) we used a uniform neutral rate of 10 substitutions per site across all genes.

Predictions using a likelihood ratio test (LRT) were performed with the BAD_Mutations pipeline (34). The pipeline is comprised of Python and Bourne Again Shell (BASH) scripts and incorporates several open-source tools, including the alignment tool PASTA (42) and maximum likelihood methods implemented in HyPhy (48). The processing step of BAD_Mutation consists of three major subcommands: (1) setup; (2) fetch; (3) align; (4) predict; and (5) compile (Figure S1). The **setup** subcommand generated the configure files. The **fetch** subcommand downloads gzipped CDS FASTA files from both Phytozome (https://phytozome.jgi.doe.gov/pz/portal.html) and Ensembl Plants (<u>http://plants.ensembl.org/index.html</u>), and then creates BLAST databases for identifying homologs. The **align** subcommand uses BLAST to identify homologs of any query protein and generates a protein alignment and phylogenetic tree using PASTA. The **predict** subcommand generates predictions for a list of codons of interest by sending a custom batch command to implement a likelihood ratio test using HyPhy. The likelihood ratio test compares the log likelihood of evolution at a single codon under a neutral model (*dN* = *dS*) to a model allowing for constraint (*dN* = *ωdS*), where *dN* and *dS* are the synonymous and nonsymous substitution rates and *ω* is a free parameter for selective constraint (Chun and Fay, 2009). The **compile** is to generate the report and p-values. The user manual, including a brief tutorial, is available at <u>https://github.com/MorrellLAB/BAD_Mutations/blob/master/Manual/Manual_v1.0.md</u>.

The BAD_Mutations pipeline makes use of sequenced and annotated genomes. We used BLAST searches of the *A. thaliana* gene sequences against 42 Angiosperm genomes, retaining the top hit from each species with a BLAST E-value threshold of 0.05. The homologue searches were restricted to Angiosperm genomes to avoid extensive saturation of synonymous sites. Protein alignments were generated with PASTA (42), and a likelihood ratio test (LRT) for constraint on each codon of interest was calculated using HyPhy (48). Sequences with ‘N’s or other ambiguous nucleotides were discarded prior to the likelihood ratio test. The LRT differs compared to its original formulation (26) in that: i) dS was estimated using all codons for each gene separately, ii) query sequences were optionally masked in the likelihood calculation to avoid any reference bias and iii) branches with *dS* greater than 3 were set to 3 to avoid spuriously high estimates of *dS*. Additionally, the original LRT used heuristics to eliminate sites with *dN* > *dS*, the derived allele present in another species, or sites with fewer than 10 species in the alignment. Rather than eliminating sites, we used logistic regression to provide a single probability of being deleterious based on the LRT test and these additional pieces of information.

Logistic regression was applied using both the masked and unmasked LRT *p*-values, where the masked *p*-values were generated from alignments without the *A. thaliana* reference allele. For the unmasked logistic regression, we used the terms log10(LRT *p*-value), *constraint* (*dN*/*dS*), *Rn*, and *An*, where *Rn* and *An* are the number of *A. thaliana* reference and alternative (i.e., mutant) amino acids observed in the alignment, respectively. For the masked model, we replaced *An* and *Rn* with the absolute value of *Rn* – *An* and the maximum of *Rn* and *An*, respectively. For both models *p*-values < 1e-16 were set to 1e-16 and constraint values > 10 were set to 10. Ten-fold cross-validation was used to assess the fit of the logistic regression. The average area under the ROC curve based on cross-validation was 0.9575 (unmasked) and 0.9471 (masked). Because these values were nearly identical to the performance of the model fit to the entire dataset, 0.9581 (unmasked) and 0.9471 (masked), we used the logistic regression coefficients from the full dataset:

> log(*p*/(1-*p*)) = −2.407-0.2139\**LRT*(*unmasked*)-0.2056\**constraint*+0.07368\**Rn*-0.1236\**An*
>
> log(*p*/(1-*p*)) = −2.453-0.1904\**LRT*(*masked*)-0.1459\**constraint*+0.2199\**max*(*Rn*,*An*)-0.2951\**abs*(*Rn*-*An*)

Sensitivity, specificity, and area under the curve (AUC) were calculated for each approach using the pROC package in R (49). We define sensitivity as the proportion of phenotype-altering variants that are predicted to be deleterious, and specificity as the proportion of variants without known phenotypic effects that are predicted to be neutral. Confidence intervals for each were calculated by 2,000 replicates of stratified bootstrapping.

Combined predictions were generated based on the combined scores of six approaches: LRT, LRT-masked, PolyPhen2, Provean, GERP++, and MAPP. SIFT 4G was not included in the combined predictions because it had missing predictions for a large number (855) variants. Sites with missing predictions from one or more of the remaining approaches (*n* = 215) were removed. Combined predictions were generated using: 1) logistic regression with each approach's score as a predictive variable, 2) support vector machine, 3) random forest, 4) linear discriminant analysis and 5) generalized linear model with penalized maximum likelihood implemented by the glmnet package in R (50). The performance of each model was assessed by AUC values obtained from 10-fold cross-validation.

## RESULTS

### Curation of a test set of *Arabidopsis thaliana* mutants

To evaluate approaches that predict deleterious variants, we generated a database of *A. thaliana* amino acid substitutions from mutants with described phenotypic alterations and common amino acid polymorphisms unlikely to affect fitness. Out of 2,910 mutants in 995 genes, 81% were from manually curated entries in UniProtKB/Swiss-Prot (*n* = 2,368), 10% were from our own literature curation (*n* = 293) and 8.6% were independently identified in both sets (*n* = 249) (Table S2). Within the same 995 genes, 1,583 common amino acid polymorphisms were identified in 80 accessions (29). For our analyses, we assume mutations that cause a deviation from the wildtype phenotype are likely deleterious.

### Performance of approaches designed to identify deleterious variants

Using the database of *A. thaliana* mutations, we assessed seven approaches for their ability to distinguish deleterious and neutral changes. The approaches were selected because they can generate predictions in non-human organisms. Comparison of sensitivity to specificity showed that each approach could reliably distinguish deleterious and neutral substitutions (Figure 1). A likelihood ratio test (LRT) implemented using the BAD_Mutations pipeline showed significantly higher performance than all other approaches as measured by the area under the curve (AUC) of sensitivity versus specificity (Figure 1, Table S3). A reference masked version of LRT (LRTm), designed to eliminate reference bias (51), was the approach with the second highest performance. PROVEAN and PolyPhen2 showed similar performance as measured by AUC, significantly higher than SIFT, GERP++ and MAPP. The relative ranking by AUC was identical when 1,050 mutations with missing predictions for at least one approach were removed (Table S3). We also found very similar measures of performance when we used common SNPs in a set of independent, randomly selected genes rather than common SNPs within the 995 genes with known phenotype altering mutations (Table S3).

**Figure 1.**
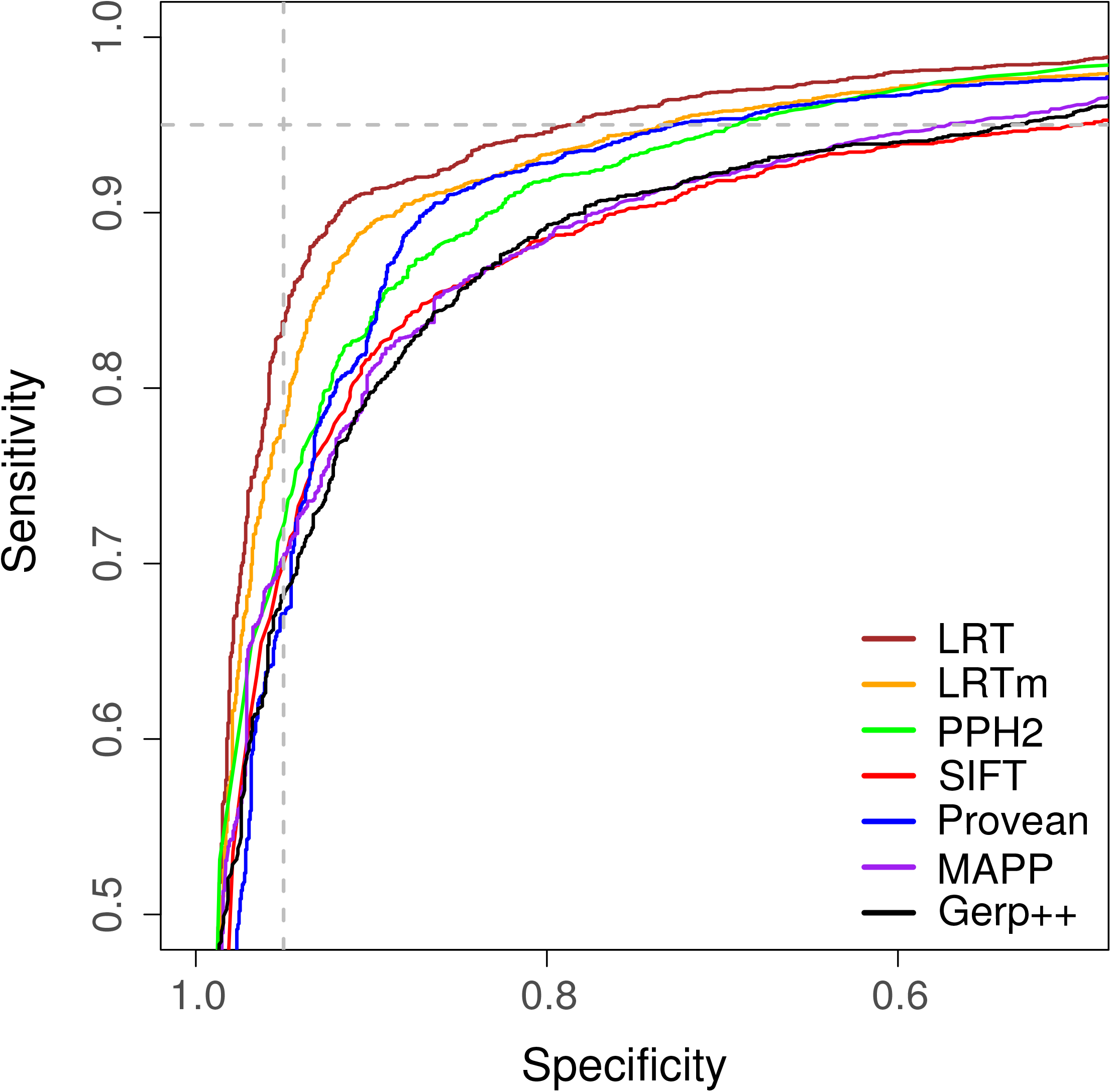
Comparison of approaches that distinguish deleterious and neutral amino acid substitutions. The fraction of true positives (sensitivity) versus the fraction of true negatives (specificity) is shown for seven approaches (LRTm is a masked version of LRT, PPH2 is PolyPhen2). The curves are based on 2,910 deleterious variants and 1,583 neutral variants. Vertical and horizontal dashed lines show the cutoff at 95% specificity and 95% sensitivity, respectively.

A second means of assessing performance is through comparing predictions of rare versus common variants. Common variants are likely neutral or nearly neutral, whereas deleterious alleles are expected to be kept at low frequency (52). Using SNPs identified in a set of 80 *A. thaliana* strains, we found each approach identified more deleterious SNPs at low compared to common frequencies (Figure 2). At minor allele frequencies between 2/80 (2.5%) and 8/80 (10%), the LRTm and SIFT predicted a lower proportion of deleterious SNPs compared to the other approaches, indicating that they are less sensitive to detecting alleles under weak selection. At the lowest frequency 1/80 (1.25%), which is expected to include many rare and potentially strongly deleterious variants, LRT called the largest proportion of SNPs deleterious.

**Figure 2.**
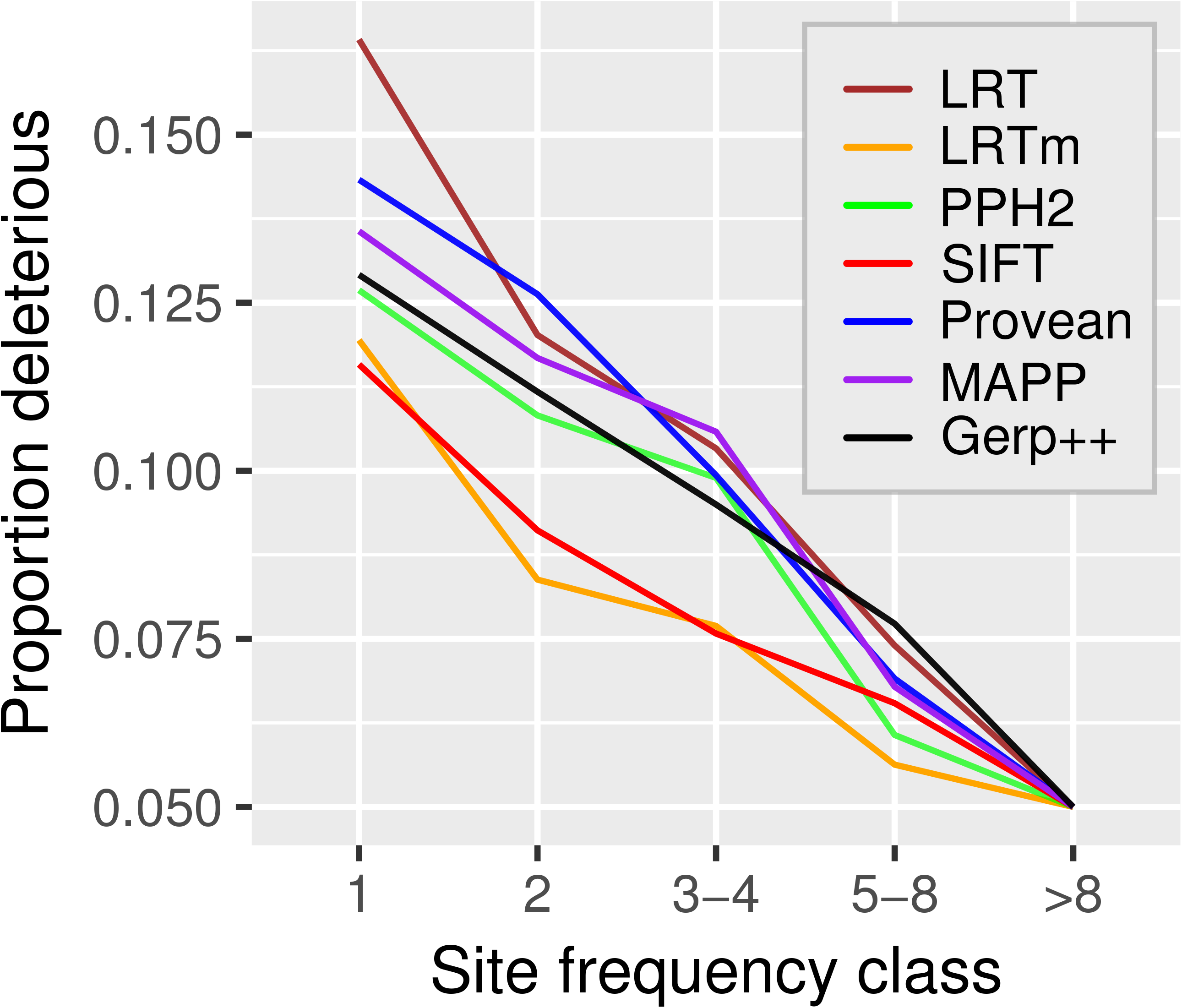
The proportion of SNPs called deleterious across frequency classes. The fraction of SNPs called deleterious by each approach (legend) at its 95% specificity threshold across five frequency classes, labeled by the number of minor alleles present (n = 80). The minor allele is defined as the allele that is less frequent in the sample. Sample sizes for the five classes are 5303 (1), 1646 (2), 1250 (3-4), 1015 (5-8) and 1583 (>8).

### Performance across phenotypic and duplicate gene categories

To further characterize differences in performance we compared class of variants, including those identified by genome-wide mutant screens or by directly targeting individual proteins. Mutants identified from screens have gross morphological or easily observable phenotypic effects and are often assigned allele names, whereas directed mutants are not often given allele names and tend to have biochemical phenotypes. To compare these two groups, we split the data into those with allele names (1,910), as a proxy for those with gross phenotypes, and those without allele names (1,000), as a proxy for biochemical phenotypes. As measured by AUC, some of the approaches performed better and their performance was more similar for the gross phenotypic class compared to the biochemical class (Figure 3a). Both SIFT and PolyPhen2 demonstrated the largest difference in performance for predicting mutations with gross phenotypic alterations versus biochemical phenotypes. For this type of mutation, the performance of PolyPhen2 was comparable to the LRT.

**Figure 3.**
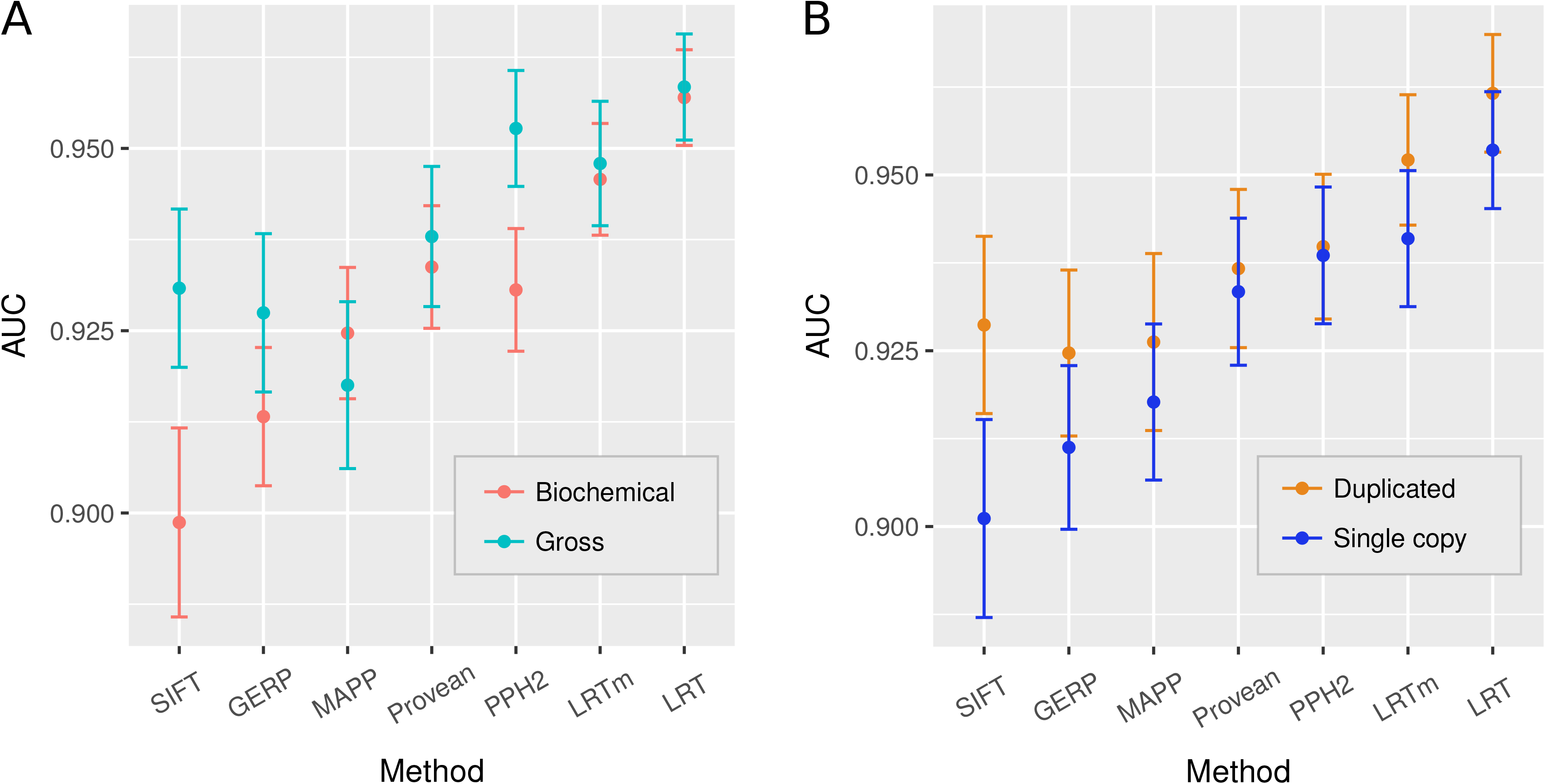
Performance of approaches across different classes of sites. Performance is measured by the area under the curve (AUC) of the approach's sensitivity versus specificity. A – comparison of mutants with biochemical (n = 1,000) versus gross phenotypes (n = 1,910). B – comparison of performance for substitutions in duplicated (n = 2,098) versus single copy genes (n = 2,395). Confidence intervals were determined by 2,000 bootstrapping iterations.

Gene duplication may reduce prior selective constraints on a protein, enabling variants to occur at previously conserved sites (53). Thus, duplicated genes may pose challenges to predicting deleterious alleles, and none of the approaches explicitly distinguish orthologs and paralogs. We identified 466 of the 995 genes as duplicated in *A. thaliana* based on BLASTP hits with 60% or more identity. We compared the performance of these genes to the remaining single copy genes. Each approach showed equal or better performance for duplicated versus single copy genes. SIFT had the largest increase in performance (Figure 3b).

### Approach dissimilarity and composite predictions

As reported previously (20, 22, 26, 54), we found substantial disagreement in predictions among the approaches. At a 95% specificity threshold, 93.6% of mutants were predicted deleterious by one or more approach but only 51.3% were predicted deleterious by at least six of the seven approaches (Table S2). Similarly, only 0.25% of common SNPs were predicted deleterious by all approaches but 16.6% were predicted deleterious by at least one approach (LRT and LRTm were considered separately). Comparing the disagreement between approaches, we found LRT and LRTm to produce very similar predictions, but to be distinct from most of the other approaches (Figure 4). We used five models that combined the predictions of all approaches except for SIFT, which had a higher proportion of missing calls. Only two of these ensemble models, a linear discriminant analysis and a generalized linear model with penalized maximum likelihood, performed significantly higher than LRT based on an AUC (Table S4).

**Figure 4.**
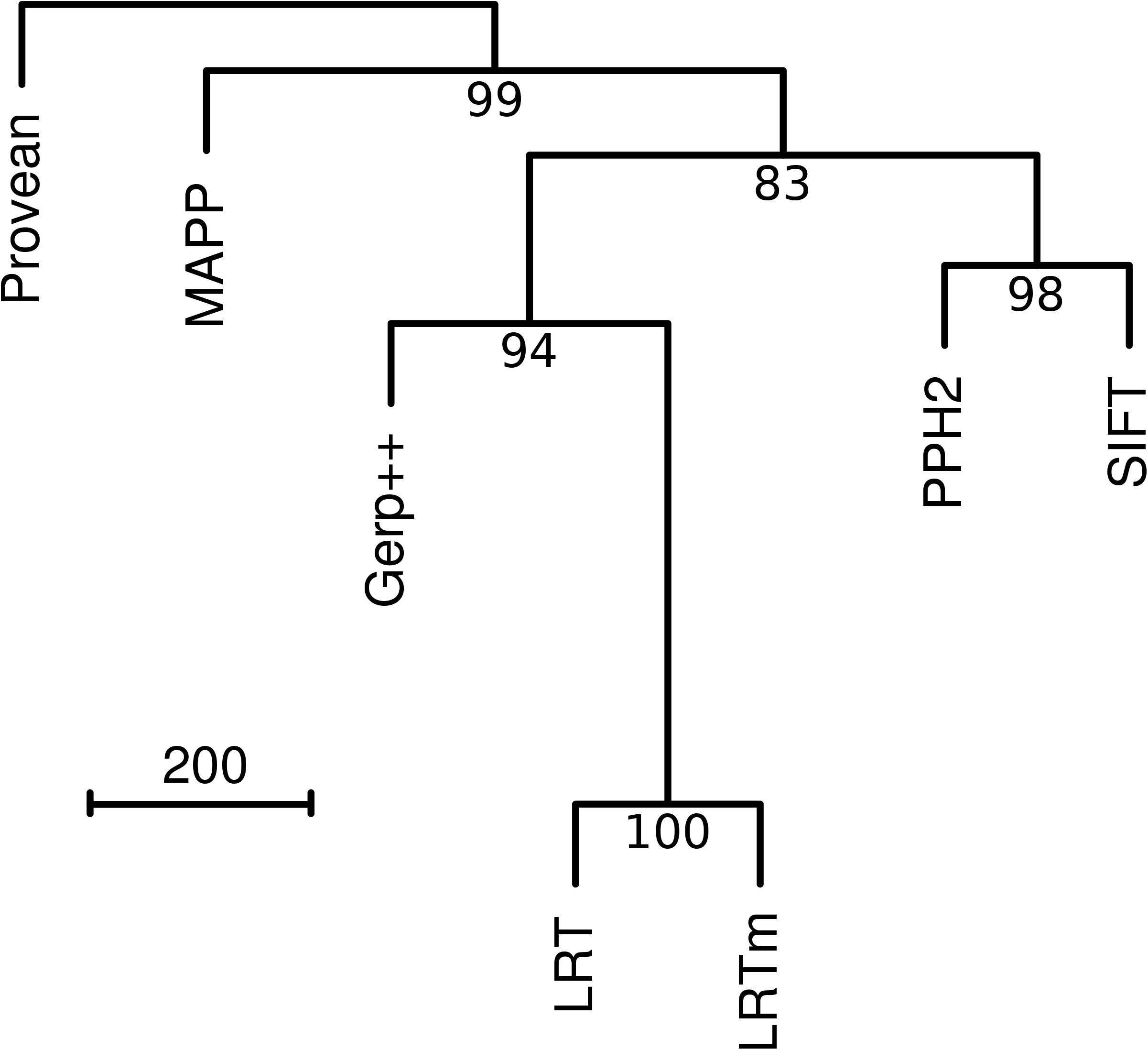
Dissimilarities among approaches. Dissimilarities were computed by the pairwise number of disagreements between each approach applied to mutants and common SNPs (n = 4,493). Dissimilarities are represented by a tree based on hierarchical clustering and values below nodes are bootstrap support based on 2,000 iterations.

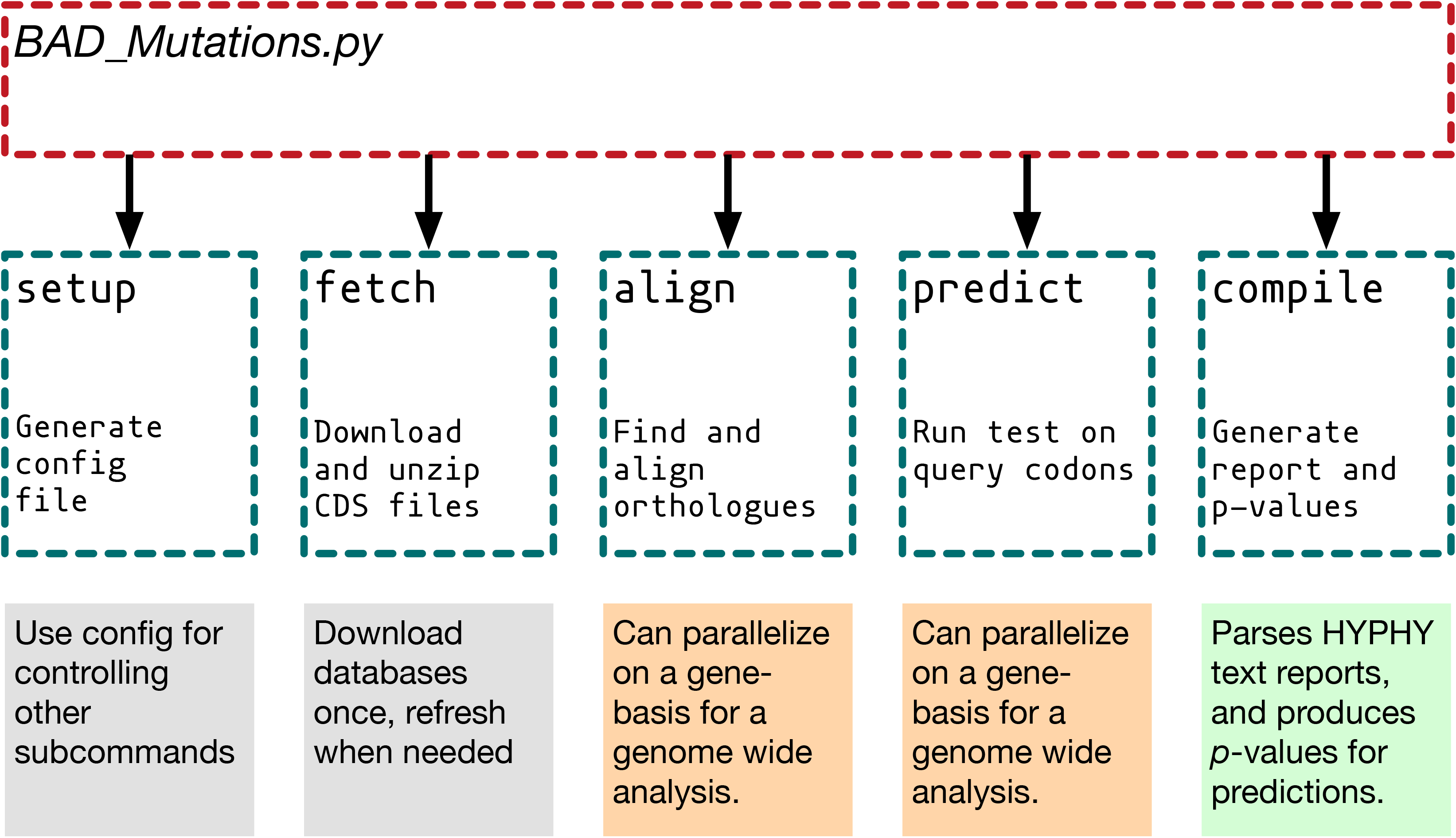

## DISCUSSION

In this study, we benchmarked the potential for several widely-used approaches to distinguish putatively deleterious and neutral amino acid substitutions in *A. thaliana*. Prior evaluations of performance focused on large sets of mutants for single proteins or known human disease variants (24, 27). Overall, we find high performance across approaches in their ability to distinguish neutral and deleterious variants, validating their use in plants. The highest performance is achieved by a likelihood ratio test (LRT) implemented using the BAD_Mutations pipeline, in this case using alignments from 42 plant genomes. However, the relative performance depended on the test set and, as discussed below, differs from previous benchmarking studies in humans. Thus, we recommend caution in interpreting slight differences in performance and advocate the use of multiple methods to achieve the highest confidence.

Below, we discuss our results along with characteristics of the approaches and test data that may contribute to differences in predictions and performance when applied to non-human species. One important consideration is the distinction between deleterious variants and those that impact protein function and have phenotypic consequences. While these two groups are overlapping, they are not identical. Because conservation between species is directly related to fitness, we have used the term “deleterious” when referring to the prediction approaches. However, the test sets used to evaluate approaches are composed of variants known to affect protein function or phenotype. Thus, regardless of the nomenclature, any evaluation of approach performance necessarily assumes a large overlap between conserved amino acid positions and those that affect protein function as measured by phenotype. Equally relevant, we use common variants as “neutral” controls even though some common variants are likely to affect protein function due to local adaptation (55) or hitchhiking (56). Despite potential contamination, common variants provide the only large set of negative controls that can be used for training and estimating rates of false positives (5). Both common and rare variants may also have compensatory effects on deleterious variants (57). These potential interactions between variants further complicates the identification of truly deleterious variants in any species.

### Phylogenetic power, alignments, and reference databases

Phylogenetic power is critical to all comparative genomic approaches that predict deleterious variants. When homologs are too closely related, not enough time has passed for neutral sites to accumulate amino acid substitutions. When homologs are too distantly related, functional sites may not be conserved due to compensatory changes or divergence in homolog function (58–60). The LRT differs from the other approaches examined in that it uses synonymous sites as an internal control to account for the expected amount of protein divergence under a neutral model. As such, even homologs that are nearly identical in their amino acid sequences are informative, given that they have accumulated changes at synonymous sites. However, distantly related homologs are uninformative if divergence at synonymous sites is saturated, thus the LRT should only be applied to organisms where a sufficient number of related genomes are available. GERP++ is similar to the LRT in that it uses a neutral substitution rate to make its predictions, but differs in that the neutral rate must be specified rather than being estimated from synonymous sites within the alignment. GERP++ also does not make use of the genetic code to distinguish synonymous and nonsynonymous changes. In this regard, GERP++ was not appropriately applied since we used a fixed neutral rate for all genes rather than an alignment specific neutral rate.

Out of the approaches compared, phylogenetic power cannot explain the differences between the LRT, MAPP, and GERP++ because they used the same alignments. However, we did notice substantial differences in performance based on the number of ungapped sequences present in the BAD_Mutations alignment at the position being queried (Figure S2). Both LRT and LRTm performed much better than the other approaches when there were 10 or fewer sequences at the position of interest. We did not see this pattern when we used the number of sequences present at any position in the alignment, which was typically close to 42. We also did not see this pattern when we examined performance based on the number of sequences used by Provean or PolyPhen2, typically over 100 per gene.

All approaches studied here use alignments to make their predictions, making the protein database and choice of homologs to be included in the alignment a critical step. For MAPP, GERP++, and LRT we used alignments generated using the BAD_Mutations pipeline which queries proteins from a set of annotated reference genomes, in this case from 42 Angiosperm species. SIFT and PolyPhen2 use the UniRef database (2011), whereas PROVEAN uses the most recent non-redundant protein database from NCBI. Both PROVEAN and PolyPhen2 are known to be sensitive to the choice of the reference database and criteria for inclusion of homologs (27, 39). Despite the choice of homologs being an important step in predicting deleterious substitutions, the use of a plant-specific or entire non-redundant database does not appear to be a major contributor to performance differences (Figure 1).

### Training and test sets

Performance of an individual approach depends on both the training and test sets used to measure it. Because performance is typically measured using common SNPs and known disease variants in humans, there has been some concern over the lack of independence between training and test sets (21, 61)(21, 61)(21, 60). However, another consideration that has not yet been examined is whether performance in one species translates to other distantly related species, which may not have the same availability of homologs from sequenced genomes spanning a range of phylogenetic relatedness. The performance of individual approaches could depend on the study system in that some approaches may expect homologs at certain phylogenetic distances, low rates of compensatory change, or low rates of gene duplication.

Previous studies of the accuracy of prediction approaches made use of five human test datasets (21, 61). We find better performance across approaches in our *A. thaliana* dataset than that reported for humans (Table 1). It is unclear why the approaches uniformly perform better in *A. thaliana*. One possibility is that the neutral and deleterious variants in *A. thaliana* are more distinct from one another than in humans. The very large proportion of phenotyping changing variants in our *A. thaliana* test set that are identified as deleterious means that this test data set is less useful for approach comparison due to the small number of cases that are difficult to predict correctly.

**Table 1.**
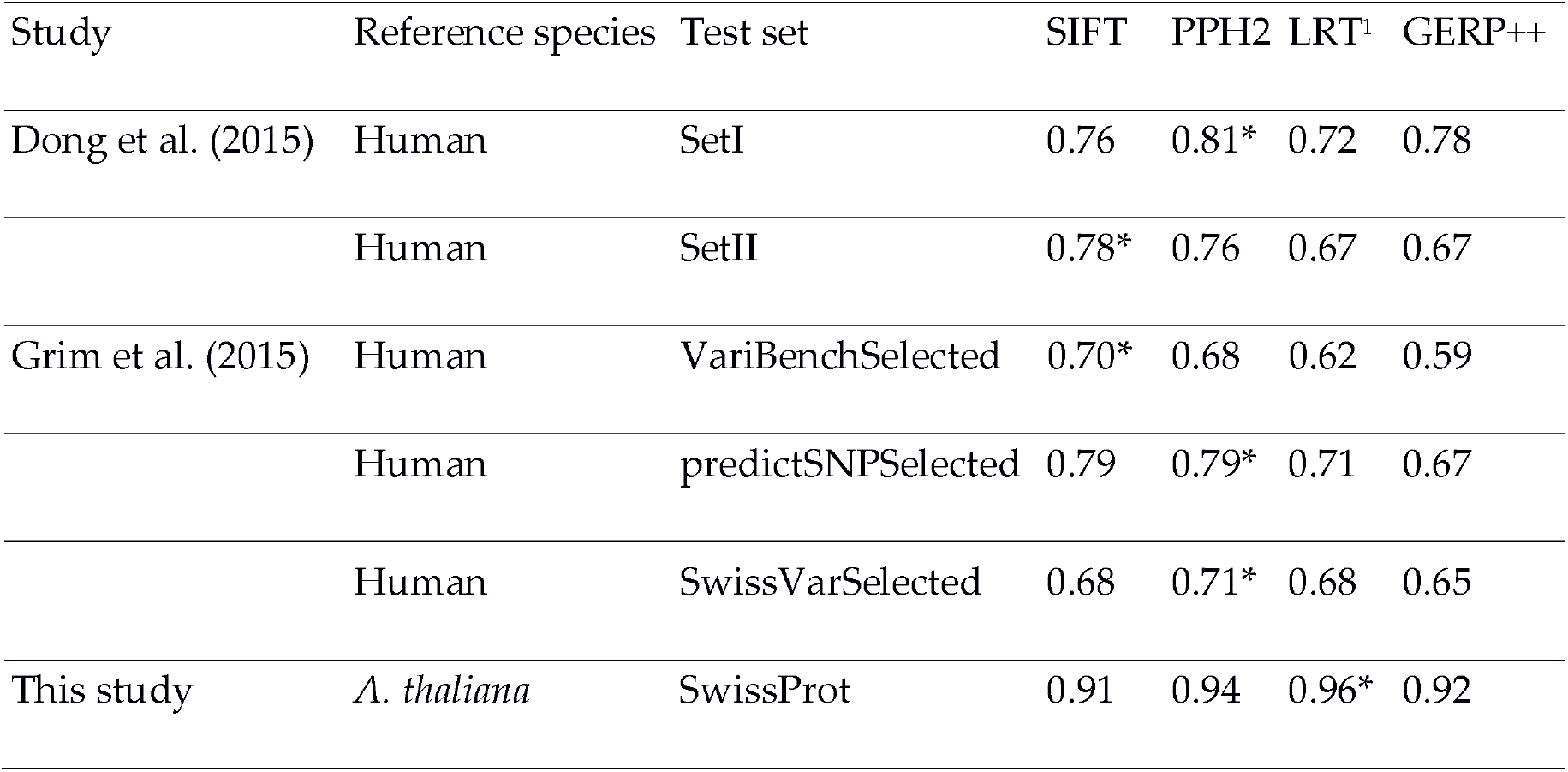
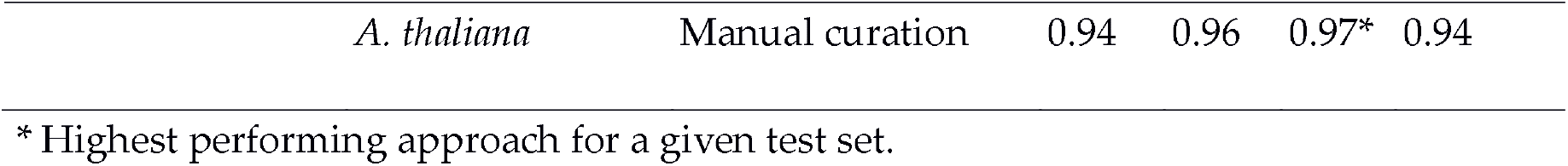
Performance measured by AUC of approaches based on different test sets.

### Population and gene-specific performance

Because nearly all measures of performance use either common polymorphism or recently fixed amino acid substitutions as a proxy for neutral SNPs, population and gene-specific factors that influence neutral polymorphism are expected to influence measures of performance. Humans have a small effective population size relative to other mammals (62) and consequently a high ratio of nonsynonymous to synonymous diversity (44, 63). Thus, distinguishing neutral and deleterious variants may be more difficult in humans than other species, and approaches trained using human polymorphism may be more conservative with respect to weakly deleterious variants. In comparison, predicting deleterious variants in *A. thaliana* may be facilitated by the fact that *A. thaliana* has slightly larger effective population size (29).

It should be noted that both demographic history and the process of local adaptation could play important roles in the distribution of deleterious variants. In populations that are colonizing or expanding into novel environments, the selective coefficients against individual variants may change (64), and locally adaptive variants may become appreciably enriched. Both humans and *A. thaliana* are known to have undergone demographic expansion in their recent evolutionary histories (65, 66). While the relative extent of local adaptation in these two species is difficult to quantify, both exhibit an excess of low-frequency amino acid polymorphism characteristic of deleterious variants (29, 67, 68).

Another potentially important factor in predicting deleterious variants is gene duplication. *A. thaliana* carries remnants of a whole genome duplication along with numerous tandem duplications (69) more than are present in the human genome (70). Gene duplication can lead to relaxed selection during subfunctionalization or pseudogenization (71), enabling amino acid variants to accumulate in recently duplicated genes. However, we found very similar performance between duplicate and single copy genes, consistent with a similar finding in humans using PolyPhen2 (27). Because we only included genes with known mutant phenotypes, the sample of recently duplicated genes is limited.

### Conclusions and future directions

Most approaches developed to predict deleterious mutations were trained using human data and in many cases, can only be used for human proteins, e.g., (7, 72, 73). This study demonstrates that several generalized approaches perform exceptionally well in *A. thaliana*, implying that they should also work well for other plant species. Because of the similarly high performance, other considerations such as ease of implementation and compute time may be considered when choosing an approach to identify deleterious mutations in plants. Notably, LRT requires longer run times than any of the other approaches, typically 5.2 hrs of computing time per gene compared to 14.5 and 9.4 minutes per gene for PolyPhen2 and Provean, respectively. One way the BAD_Mutations pipeline could be sped up while retaining the flexibility of querying customizable plant genomes is by using heuristic measures of site-specific conservation rather than the LRT. Provocatively, we found similar performance (AUC = 0.9551) for a logistic model that only used the number of reference and alternative alleles in the alignment (*Rn* and *An*). However, such heuristic measures may not be robust to a change in the reference species and its distance to other genomes in the database. A second approach would be to use predictions from the combined output of multiple prediction approaches, as this has been shown to be highly effective in humans, e.g., (20). Although we did not find an ensemble predictor that greatly improved performance, removing LRT predictions did not reduce the performance of the ensemble predictions.

## AVAILABILITY

LRT predictions were implemented in the Python package BAD_Mutations which is freely available from http://github.com/MorrellLAB/BAD_Mutations.git.

## ACKNOWLEDGEMENTS

We thank members of the Morrell Lab for discussion and software testing. We also would like to thank Dr. Danelle Seymour for helpful comments on an earlier version of the manuscript. Hardware and software support were provided by the University of Minnesota Supercomputing Institute.

## Author contributions

T.J.Y.K. and P.J.H. wrote code for BAD_Mutations. J.C.F., L. L. and C. H. S. analyzed the data. L.L., J.C.F., T.J.Y.K., and P.L.M. wrote the initial draft of the manuscript. All authors contributed to final manuscript preparation.

---

Table S1. Summary of different methods used to predict deleterious variants.

Table S2. Amino acid substitutions used to measure performance and predictions for seven approaches. The amino acid substitutions include 2,910 with phenotypic effects and 10,797 SNPs in 995 *A. thaliana* genes. A list of 2,617 amino acid-altering mutations in 960 *A. thaliana* genes

Table S3. Performance of methods used to distinguish deleterious and neutral substitutions Table S4. Performance of models based on ensemble prediction methods.

Figure S1. Overview of the BAD_Mutations pipeline. BAD_Mutations consists of three sequential commands: fetch, align and predict. The fetch command downloads CDS FASTA files from the Phytozome and Ensembl Plant databases and converts them into BLAST databases. The align command runs BLAST to identify homologs and PASTA to generate multiple sequence alignments and phylogenetic trees based on the protein sequence. The predict command runs a HyPhy script to generate predictions for a list of input substitutions.

Figure S2. Performance depends on the number of aligned sequences. Performance was measured by the area under the curve (AUC) of sensitivity versus specificity (lines) and 95% confidence intervals were obtained from 2,000 bootstraps (shaded areas). The number of ungapped sequences at the amino acid position of interest was obtained from BAD_Mutations alignments of proteins from 42 plant genomes. AUC was calculated for four groups binned by the number of aligned sequences.

## REFERENCES

1. Ahituv,N., Kavaslar,N., Schackwitz,W., Ustaszewska,A., Martin,J., Hébert,S., Doelle,H., Ersoy,B., Kryukov,G., Schmidt,S., et al. (2007) Medical sequencing at the extremes of human body mass. Am. J. Hum. Genet., 80, 779–791.

2. Cooper,G.M. and Shendure,J. (2011) Needles in stacks of needles: finding disease-causal variants in a wealth of genomic data. Nat. Rev. Genet., 12, 628–640.

3. Manolio,T.A., Collins,F.S., Cox,N.J., Goldstein,D.B., Hindorff,L.A., Hunter,D.J., McCarthy,M.I., Ramos,E.M., Cardon,L.R., Chakravarti,A., et al. (2009) Finding the missing heritability of complex diseases. Nature, 461, 747–753.

4. Thornton,K.R., Foran,A.J. and Long,A.D. (2013) Properties and modeling of GWAS when complex disease risk is due to non-complementing, deleterious mutations in genes of large effect. PLOS Genet., 9, e1003258.

5. Ng,P.C. and Henikoff,S. (2006) Predicting the effects of amino acid substitutions on protein function. Annu. Rev. Genomics Hum. Genet., 7, 61–80.

6. Pollard,K.S., Hubisz,M.J., Rosenbloom,K.R. and Siepel,A. (2010) Detection of nonneutral substitution rates on mammalian phylogenies. Genome Res., 20, 110–121.

7. Kircher,M., Witten,D.M., Jain,P., O’Roak,B.J., Cooper,G.M. and Shendure,J. (2014) A general framework for estimating the relative pathogenicity of human genetic variants. Nat. Genet., 46, 310–315.

8. 1000 Genomes Project Consortium,T. 1000 G.P. (2012) An integrated map of genetic variation from 1,092 human genomes. Nature, 491, 56–65.

9. Fay,J.C. (2013) The molecular basis of phenotypic variation in yeast. Curr. Opin. Genet. Dev., 23, 672–677.

10. Stenson,P.D., Mort,M., Ball,E.V., Shaw,K., Phillips,A.D. and Cooper,D.N. (2014) The human gene mutation database: building a comprehensive mutation repository for clinical and molecular genetics, diagnostic testing and personalized genomic medicine. Hum. Genet., 133, 1–9.

11. Dudley,J.T., Chen,R., Sanderford,M., Butte,A.J. and Kumar,S. (2012) Evolutionary meta-analysis of association studies reveals ancient constraints affecting disease marker discovery. Mol. Biol. Evol., 29, 2087–2094.

12. Charlesworth,D. and Willis,J.H. (2009) The genetics of inbreeding depression. Nat. Rev. Genet., 10, 783–796.

13. Yang,J., Mezmouk,S., Baumgarten,A., Buckler,E.S., Guill,K.E., McMullen,M.D., Mumm,R.H. and Ross-Ibarra,J. (2017) Incomplete dominance of deleterious alleles contributes substantially to trait variation and heterosis in maize. PLOS Genet., 13, e1007019.

14. Cruz,F., Vilà,C. and Webster,M.T. (2008) The legacy of domestication: accumulation of deleterious mutations in the dog genome. Mol. Biol. Evol., 25, 2331–2336.

15. Liu,Q., Zhou,Y., Morrell,P.L. and Gaut,B.S. (2017) Deleterious variants in Asian rice and the potential cost of domestication. Mol. Biol. Evol., 10.1093/molbev/msw296.

16. Lu,J., Tang,T., Tang,H., Huang,J., Shi,S. and Wu,C.-I. (2006) The accumulation of deleterious mutations in rice genomes: a hypothesis on the cost of domestication. Trends Genet., 22, 126–131.

17. Rodgers-Melnick,E., Bradbury,P.J., Elshire,R.J., Glaubitz,J.C., Acharya,C.B., Mitchell,S.E., Li,C., Li,Y. and Buckler,E.S. (2015) Recombination in diverse maize is stable, predictable, and associated with genetic load. Proc. Natl. Acad. Sci., 112, 3823–3828.

18. Moyers,B.T., Morrell,P.L. and McKay,J.K. Genetic costs of domestication and improvement. J. Hered., 10.1093/jhered/esx069.

19. Morrell,P.L., Buckler,E.S. and Ross-Ibarra,J. (2012) Crop genomics: advances and applications. Nat. Rev. Genet., 13, 85–96.

20. González-Pérez,A. and López-Bigas,N. (2011) Improving the assessment of the outcome of nonsynonymous SNVs with a consensus deleteriousness score, condel. Am. J. Hum. Genet., 88, 440–449.

21. Grimm,D.G., Azencott,C.-A., Aicheler,F., Gieraths,U., MacArthur,D.G., Samocha,K.E., Cooper,D.N., Stenson,P.D., Daly,M.J., Smoller,J.W., et al. (2015) The evaluation of tools used to predict the impact of missense variants is hindered by two types of circularity. Hum. Mutat., 36, 513–523.

22. Olatubosun,A., Väliaho,J., Härkönen,J., Thusberg,J. and Vihinen,M. (2012) PON-P: integrated predictor for pathogenicity of missense variants. Hum. Mutat., 33, 1166–1174.

23. Thusberg,J., Olatubosun,A. and Vihinen,M. (2011) Performance of mutation pathogenicity prediction methods on missense variants. Hum. Mutat., 32, 358–368.

24. Ng,P.C. and Henikoff,S. (2003) SIFT: predicting amino acid changes that affect protein function. Nucleic Acids Res., 31, 3812–3814.

25. Davydov,E.V., Goode,D.L., Sirota,M., Cooper,G.M., Sidow,A. and Batzoglou,S. (2010) Identifying a high fraction of the human genome to be under selective constraint using GERP++. PLOS Comput Biol, 6, e1001025.

26. Chun,S. and Fay,J.C. (2009) Identification of deleterious mutations within three human genomes. Genome Res., 19, 1553–1561.

27. Adzhubei,I., Jordan,D.M. and Sunyaev,S.R. (2013) Predicting functional effect of human missense mutations using PolyPhen-2. Curr. Protoc. Hum. Genet. Editor. Board Jonathan Haines Al, 0 7, Unit7.20.

28. Hicks,S., Wheeler,D.A., Plon,S.E. and Kimmel,M. (2011) Prediction of missense mutation functionality depends on both the algorithm and sequence alignment employed. Hum. Mutat., 32, 661–668.

29. Cao,J., Schneeberger,K., Ossowski,S., Günther,T., Bender,S., Fitz,J., Koenig,D., Lanz,C., Stegle,O., Lippert,C., et al. (2011) Whole-genome sequencing of multiple *Arabidopsis thaliana* populations. Nat. Genet., 43, 956–963.

30. Günther,T. and Schmid,K.J. (2010) Deleterious amino acid polymorphisms in *Arabidopsis thaliana* and rice. Theor. Appl. Genet., 121, 157–168.

31. Mezmouk,S. and Ross-Ibarra,J. (2014) The pattern and distribution of deleterious mutations in maize. G3 GenesGenomesGenetics, 4, 163–171.

32. Renaut,S. and Rieseberg,L.H. (2015) The accumulation of deleterious mutations as a consequence of domestication and improvement in sunflowers and other compositae crops. Mol. Biol. Evol., 32, 2273–2283.

33. Zhang,M., Zhou,L., Bawa,R., Suren,H. and Holliday,J.A. (2016) Recombination rate variation, hitchhiking, and demographic history shape deleterious load in poplar. Mol. Biol. Evol., 10.1093/molbev/msw169.

34. Kono,T.J.Y., Fu,F., Mohammadi,M., Hoffman,P.J., Liu,C., Stupar,R.M., Smith,K.P., Tiffin,P., Fay,J.C. and Morrell,P.L. (2016) The role of deleterious substitutions in crop genomes. Mol. Biol. Evol., 33, 2307–2317.

35. Goodstein,D.M., Shu,S., Howson,R., Neupane,R., Hayes,R.D., Fazo,J., Mitros,T., Dirks,W., Hellsten,U., Putnam,N., et al. (2012) Phytozome: a comparative platform for green plant genomics. Nucleic Acids Res., 40, D1178–D1186.

36. Lockton,S. and Gaut,B.S. (2005) Plant conserved non-coding sequences and paralogue evolution. Trends Genet., 21, 60–65.

37. Comai,L. (2005) The advantages and disadvantages of being polyploid. Nat. Rev. Genet., 6, nrg1711.

38. Charlesworth,B. (2012) The role of background selection in shaping patterns of molecular evolution and variation: evidence from variability on the *Drosophila* X chromosome. Genetics, 191, 233–246.

39. Choi,Y., Sims,G.E., Murphy,S., Miller,J.R. and Chan,A.P. (2012) Predicting the functional effect of amino acid substitutions and indels. PLOS ONE, 7, e46688.

40. Stone,E.A. and Sidow,A. (2005) Physicochemical constraint violation by missense substitutions mediates impairment of protein function and disease severity. Genome Res., 15, 978–986.

41. Altschul,S.F., Gish,W., Miller,W., Myers,E.W. and Lipman,D.J. (1990) Basic local alignment search tool. J. Mol. Biol., 215, 403–410.

42. Mirarab,S., Nguyen,N., Guo,S., Wang,L.-S., Kim,J. and Warnow,T. (2015) PASTA: ultra-large multiple sequence alignment for nucleotide and amino-acid sequences. J. Comput. Biol., 22, 377–386.

43. Boutet,E., Lieberherr,D., Tognolli,M., Schneider,M., Bansal,P., Bridge,A.J., Poux,S., Bougueleret,L. and Xenarios,I. (2016) UniProtKB/Swiss-Prot, the manually annotated section of the UniProt KnowledgeBase: how to use the entry view. In Edwards,D. (ed). Plant Bioinformatics. Springer New York, New York, NY, Vol. 1374, pp. 23–54.

44. Fay,J.C., Wyckoff,G.J. and Wu,C.-I. (2001) Positive and negative selection on the human genome. Genetics, 158, 1227–1234.

45. Boyko,A.R., Williamson,S.H., Indap,A.R., Degenhardt,J.D., Hernandez,R.D., Lohmueller,K.E., Adams,M.D., Schmidt,S., Sninsky,J.J., Sunyaev,S.R., et al. (2008) Assessing the evolutionary impact of amino acid mutations in the human genome. PLOS Genet., 4, e1000083.

46. Adzhubei,I.A., Schmidt,S., Peshkin,L., Ramensky,V.E., Gerasimova,A., Bork,P., Kondrashov,A.S. and Sunyaev,S.R. (2010) A method and server for predicting damaging missense mutations. Nat. Methods, 7, 248–249.

47. Vaser,R., Adusumalli,S., Leng,S.N., Sikic,M. and Ng,P.C. (2016) SIFT missense predictions for genomes. Nat. Protoc., 11, 1–9.

48. Pond,S.L.K., Frost,S.D.W. and Muse,S.V. (2005) HyPhy: hypothesis testing using phylogenies. Bioinformatics, 21, 676–679.

49. Robin,X., Turck,N., Hainard,A., Tiberti,N., Lisacek,F., Sanchez,J.-C. and Müller,M. (2011) pROC: an open-source package for R and S+ to analyze and compare ROC curves. BMC Bioinformatics, 12, 77.

50. Friedman,J., Hastie,T. and Tibshirani,R. (2010) Regularization paths for generalized linear models via coordinate descent. J. Stat. Softw., 33, 1–22.

51. Simons,Y.B., Turchin,M.C., Pritchard,J.K. and Sella,G. (2014) The deleterious mutation load is insensitive to recent population history. Nat. Genet., 46, 220–224.

52. Ewens,W.J. (2004) Mathematical population genetics Springer New York, New York, NY.

53. Kondrashov,F.A., Rogozin,I.B., Wolf,Y.I. and Koonin,E.V. (2002) Selection in the evolution of gene duplications. Genome Biol., 3, research0008.

54. Doniger,S.W., Kim,H.S., Swain,D., Corcuera,D., Williams,M., Yang,S.-P. and Fay,J.C. (2008) A catalog of neutral and deleterious polymorphism in yeast. PLOS Genet., 4, e1000183.

55. Hancock,A.M., Brachi,B., Faure,N., Horton,M.W., Jarymowycz,L.B., Sperone,F.G., Toomajian,C., Roux,F. and Bergelson,J. (2011) Adaptation to climate across the *Arabidopsis thaliana* genome. Science, 334, 83–86.

56. Chun,S. and Fay,J.C. (2011) Evidence for hitchhiking of deleterious mutations within the human genome. PLOS Genet., 7, e1002240.

57. Poon,A., Davis,B.H. and Chao,L. (2005) The coupon collector and the suppressor mutation: estimating the number of compensatory mutations by maximum likelihood. Genetics, 170, 1323–1332.

58. Breen,M.S., Kemena,C., Vlasov,P.K., Notredame,C. and Kondrashov,F.A. (2012) Epistasis as the primary factor in molecular evolution. Nature, 490, 535–538.

59. Jordan,D.M., Frangakis,S.G., Golzio,C., Cassa,C.A., Kurtzberg,J., Task Force for Neonatal Genomics, Davis,E.E., Sunyaev,S.R. and Katsanis,N. (2015) Identification of cis-suppression of human disease mutations by comparative genomics. Nature, 524, 225–229.

60. Marini,N.J., Thomas,P.D. and Rine,J. (2010) The use of orthologous sequences to predict the impact of amino acid substitutions on protein function. PLOS Genet., 6, e1000968.

61. Dong,C., Wei,P., Jian,X., Gibbs,R., Boerwinkle,E., Wang,K. and Liu,X. (2015) Comparison and integration of deleteriousness prediction methods for nonsynonymous SNVs in whole exome sequencing studies. Hum. Mol. Genet., 24, 2125–2137.

62. Leffler,E.M., Bullaughey,K., Matute,D.R., Meyer,W.K., Ségurel,L., Venkat,A., Andolfatto,P. and Przeworski,M. (2012) Revisiting an old riddle: what determines genetic diversity levels within species. PLOS Biol., 10, e1001388.

63. Kosiol,C., Vina•,T., Fonseca,R.R. da, Hubisz,M.J., Bustamante,C.D., Nielsen,R. and Siepel,A. (2008) Patterns of positive selection in six mammalian genomes. PLOS Genet., 4, e1000144.

64. Slotte,T., Hazzouri,K.M., Ågren,J.A., Koenig,D., Maumus,F., Guo,Y.-L., Steige,K., Platts,A.E., Escobar,J.S., Newman,L.K., et al. (2013) The *Capsella rubella* genome and the genomic consequences of rapid mating system evolution. Nat. Genet., 45, 831–835.

65. Hoffmann,M.H. (2002) Biogeography of *Arabidopsis thaliana* L. Heynh. (Brassicaceae). J. Biogeogr., 29, 125–134.

66. Finlayson,C. (2005) Biogeography and evolution of the genus *Homo*. Trends Ecol. Evol., 20, 457–463.

67. Lohmueller,K.E., Indap,A.R., Schmidt,S., Boyko,A.R., Hernandez,R.D., Hubisz,M.J., Sninsky,J.J., White,T.J., Sunyaev,S.R., Nielsen,R., et al. (2008) Proportionally more deleterious genetic variation in European than in African populations. Nature, 451, 994–997.

68. Henn,B.M., Botigué,L.R., Peischl,S., Dupanloup,I., Lipatov,M., Maples,B.K., Martin,A.R., Musharoff,S., Cann,H., Snyder,M.P., et al. (2016) Distance from sub-Saharan Africa predicts mutational load in diverse human genomes. Proc. Natl. Acad. Sci., 113, E440–E449.

69. The *Arabidopsis* Genome Initiative (2000) Analysis of the genome sequence of the flowering plant *Arabidopsis thaliana*. Nature, 408, 796–815.

70. Lynch,M. and Conery,J.S. (2000) The evolutionary fate and consequences of duplicate genes. Science, 290, 1151–1155.

71. Ohno,S. (1970) Evolution by gene duplication Springer Berlin Heidelberg, Berlin, Heidelberg.

72. Li,B., Krishnan,V.G., Mort,M.E., Xin,F., Kamati,K.K., Cooper,D.N., Mooney,S.D. and Radivojac,P. (2009) Automated inference of molecular mechanisms of disease from amino acid substitutions. Bioinformatics, 25, 2744–2750.

73. Schwarz,J.M., Rödelsperger,C., Schuelke,M. and Seelow,D. (2010) MutationTaster evaluates disease-causing potential of sequence alterations. Nat. Methods, 7, 575–576.

